# A cell-type specific surveillance complex represses cryptic promoters during differentiation in an adult stem cell lineage

**DOI:** 10.1101/2025.02.25.640250

**Authors:** Neuza R. Matias, Lorenzo Gallicchio, Dan Lu, Jongmin J. Kim, Julian Perez, Angela M. Detweiler, Chenggang Lu, Benjamin Bolival, Margaret T. Fuller

## Abstract

Regulators of chromatin accessibility play key roles in cell fate transitions, triggering onset of novel transcription programs as cells differentiate. In the *Drosophila* male germ line stem cell lineage, tMAC, a master regulator of spermatocyte differentiation that binds thousands of loci, is required for local opening of chromatin, allowing activation of spermatocyte-specific promoters. Here we show that a cell-type specific surveillance system involving the multiple zinc finger protein Kmg and the pipsqueak domain protein Dany dampens transcriptional output from weak tMAC dependent promoters and blocks tMAC binding at thousands of additional cryptic promoters, thus preventing massive expression of aberrant protein-coding transcripts. ChIP-seq showed Kmg enriched at the tMAC-bound promoters it repressed, consistent with direct action. In contrast, Kmg and Dany did not repress highly expressed tMAC dependent genes, where they colocalized with their binding partner, the chromatin modeler Mi-2 (NuRD), along the transcribed regions rather than at the promoter. Mi-2 has been shown to preferentially bind RNA over chromatin (Ullah *et al*. 2022). We propose that at highly expressed genes binding of Mi-2 to the abundant nascent RNA pulls the Kmg/Dany complex away from promoters, providing a mechanism to effectively repress ectopic promoters while protecting robust transcription.

## Introduction

During both embryonic development and cellular differentiation, pioneer transcription factors play key roles in cell state transitions due to their ability to convert previously silent chromatin from closed in precursor cells to open for productive expression of new, cell-type specific transcription program(s). Pioneer transcription factors can scan chromatin, recognize and bind target DNA sequences wrapped around nucleosomes, and locally alter the structural relationship between nucleosome, DNA, and nearby H1 linker histones, making local genomic regions accessible to downstream effectors of transcription (reviewed in (Barral and Zaret 2024)). However, the ability of pioneer factors to recognize target sites in closed chromatin raises two outstanding questions. First, given that these relatively short DNA motifs occur frequently in the genome, what mechanisms direct and restrict pioneer factor binding and action to the appropriate enhancer and promoter regions? Second, once pioneer factor binding has initiated opening of local chromatin, what mechanisms regulate whether transcription indeed initiates at those sites? Some pioneer factors have been shown to bind many genomic sites that do not become transcriptionally active (reviewed in (MacQuarrie *et al*. 2011)). For example MyoD, a pioneer factor that orchestrates differentiation of skeletal muscle, binds 30,000– 60,000 sites in different muscle cell types, the majority of which did not overlap with genes regulated by MyoD (Cao *et al*. 2010).

Analysis of how a novel robust transcription program turns on as male germ cells differentiate from proliferating spermatogonia into spermatocytes in *Drosophila* has uncovered a cell-type specific master regulatory complex required to open chromatin at thousands of promoters for spermatocyte-specific transcription, as well as a surveillance mechanism that restrains this action at cryptic promoters (Figure 1A). In the *Drosophila* male germ line stem cell lineage, transit amplifying spermatogonia stop dividing, complete a final S phase, and enter meiotic prophase as spermatocytes. During this stage of differentiation, over 1800 genes are strongly upregulated, many of which are only expressed in spermatocytes. In addition, another ∼1200 genes expressed in spermatogonia become transcribed from an alternative, spermatocyte-specific promotor (Lu *et al*.. 2020). Genetic analysis revealed that much of this novel, spermatocyte-specific transcription program requires promotor proximal action of tMAC, a cell-type specific protein complex that shows many of the hallmarks of a pioneer transcription factor, although this has not yet been assessed by biochemical tests.

**Figure 1:**
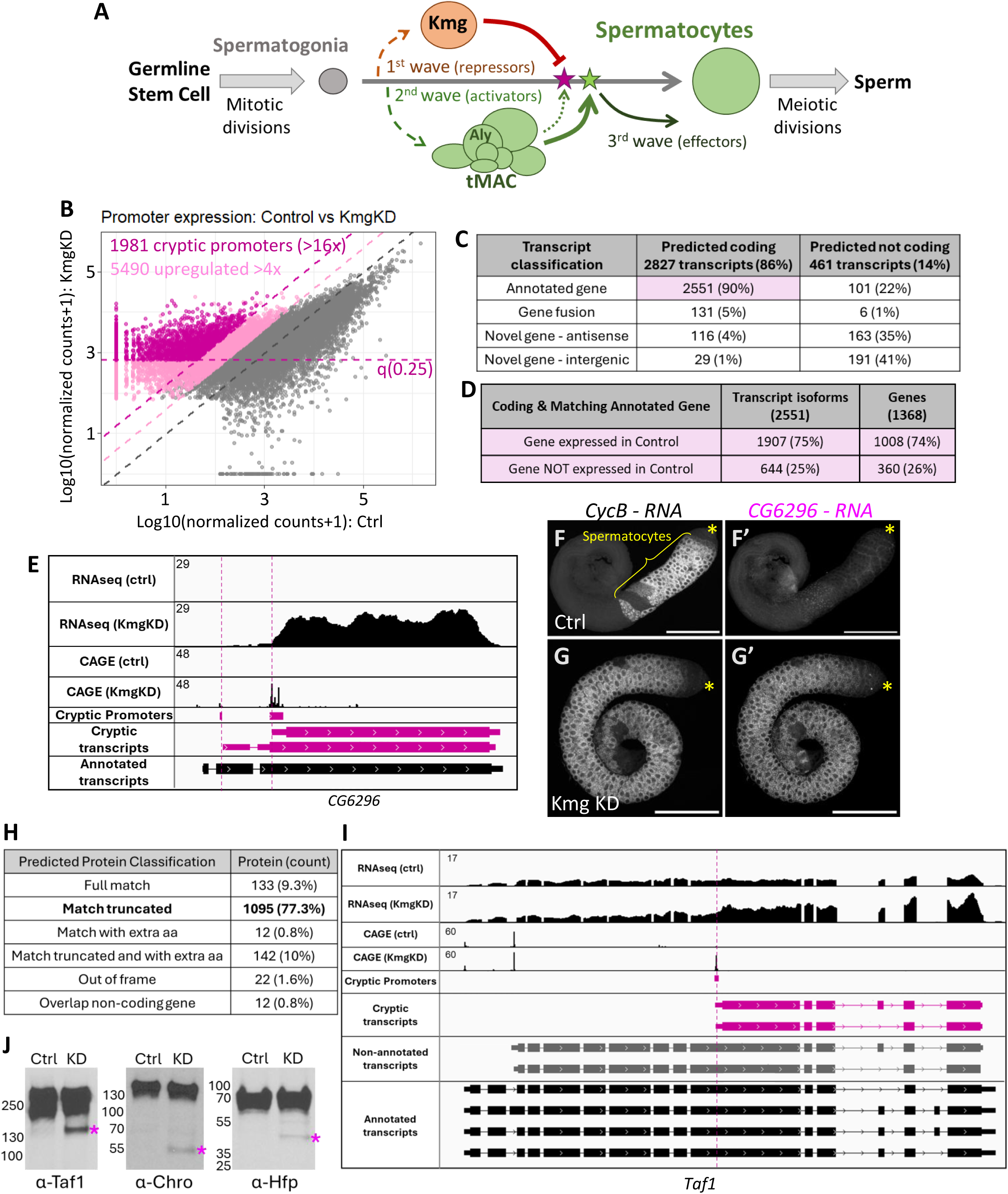
Kmg represses thousands of cryptic promoters genome wide. (A) Diagram of *Drosophila* spermatogenesis featuring the transcriptional waves after spermatogonia differentiate into spermatocytes. (B) Expression profile for all promoters expressed in testes from control [X-axis] and Kmg knockdown [Y axis]. Axes read counts normalized by DESeq2, with log10 transformation. Light pink: promoters expressed >4 fold in Kmg knockdown compared to control. Dark pink: cryptic promoters, definition in methods session. (C) Classification of cryptic transcripts in relation to annotated transcripts, by SQANTI3. (D) Classification of cryptic transcripts that matched annotated genes and are predicted to code proteins in relation to whether the host gene is expressed higher than quantile 0.2 in control testes. (E) RNA-seq and CAGE profiles for *CG6296* gene. Pink: cryptic promoters and resulting transcripts. (F-G) HCR-FISH using probes against CycB or cryptic CG6296 RNA, in control (F, F’) or Kmg knockdown (G,G’) testes. Asterisk: testis apical tip. Scale bars, 200 µm. (H) Classification of predicted cryptic proteins in relation to the most similar annotated isoform of the host gene. (I) RNA-seq and CAGE profiles for Taf1 gene. Pink: cryptic promoters and resulting transcripts. Grey: transcripts detected in control testes, but not previously annotated in *Drosophila* genome. (J) Western blots of control or Kmg knockdown testis lysates probed with antibodies against Taf1, Chro or Hfp. Due to similar size, the two Taf1 predicted cryptic proteins would not have been resolved.

tMAC is a spermatocyte-specific version of the conserved, generally expressed MuvB complex (MMB/DREAM) (Beall *et al*. 2007), which works with a variety of partners such as E2F, Rb or Myb to repress and/or activate key cell cycle and developmental genes (Sadasivam and DeCaprio 2013; Fischer and Müller 2017). The MuvB complex has been shown to bind and remodel nucleosomes, exposing nucleosomal DNA, consistent with function as a pioneer transcription complex (Koliopoulos *et al*. 2022). The spermatocyte-specific tMAC complex has two subunits also in MuvB complex (p55-Caf1 and Mip40 [LIN37]), but has replaced several other MuvB subunits with paralogs expressed only in spermatocytes. For example, in place of Mip130 [LIN9], the tMAC complex contains Aly, a Mip130 paralog first expressed in early spermatocytes (White-Cooper *et al*. 2000; Jiang and White-Cooper 2003; Beall *et al*. 2007; Jiang *et al*. 2007; Doggett *et al*. 2011).

Function of tMAC is required to turn on a following wave of spermatocyte-specific transcription that endows the male germ cells with hundreds of gene products needed for the meiotic cell cycle and post-meiotic spermatid differentiation (Figure 1A). ChIP-seq for Aly indicated binding in a peak near target promoters, with action of Aly required for a small stretch of local chromatin to convert from closed in spermatogonia to open and accessible to ATAC-seq in spermatocytes (Lu *et al*. 2020). Expression from many tMAC dependent promoters is upregulated by action of Mediator and the tTAFs, spermatocyte-specific homologs of components of TFIID (C. Lu and Fuller 2015; M. A. Hiller *et al*. 2001; M. Hiller *et al*. 2004)

We have discovered a developmentally controlled genetic regulatory system that keeps the powerful but potentially promiscuous tMAC complex from turning on transcription from thousands of inappropriate promoters across the genome, while allowing robust transcription from the normal slate of Aly-dependent spermatocyte-specific promoters. We had previously shown that a spermatocyte-specific, multiple zinc finger protein, Kumgang (Kmg), first expressed in very early spermatocytes, is required to prevent the tMAC component Aly from binding to and activating cryptic promoters associated with ∼400 genes (Kim *et al*. 2017). Because changes in expression were assessed by microarray, with probe sets only at the 3’ ends of then annotated genes, Kim *et al*. (2017) focused on loss of Kmg function causing misexpression of genes normally expressed in other tissues, but not male germ cells. Surprisingly, ChIP-seq from wild type testes showed Kmg protein localized along the gene bodies of highly expressed genes, rather than enriched at the cryptic promoters (Kim *et al*. 2017), leaving the mechanism by which *Kmg* functions to repress transcription from cryptic promoters mysterious. Kim *et al*. (2017) suggested that Kmg might prevent expression from cryptic promoters indirectly, by concentrating tMAC at its normally bound promoters, thus limiting tMAC availability to bind and activate cryptic promoters.

Here we present new data that suggest Kmg acts locally at Aly-bound sites to dampen transcription from weak promoters and, with partner proteins, also acts to repress transcription, close chromatin and reduce tMAC binding at cryptic promoters. We identified structural binding partners of Kmg by immunoprecipitation and mass spectrometry (IP-MS), including Dany, a spermatocyte-specific pipsqueak-domain protein also required for normal transcription in spermatocytes (Trost *et al*. 2016), the chromatin remodeler Mi-2 (also identified by Kim *et al.,* 2017), and the NuRD complex component Simj. Using a promoter centric analysis combining full length Pac-Bio RNA sequencing (Iso-seq), short-read RNA-seq, and CAGE (Cap Analysis Gene Expression, which identifies transcription start sites), we found that loss of either Kmg, Dany, Mi-2 or Simj function in spermatocytes upregulated expression from thousands of cryptic promoters not annotated as utilized in testes or any other *Drosophila* tissues, resulting in expression of many abnormal transcripts predicted to encode proteins that are truncated, have altered N-terminal domains, or are expressed in the wrong cell type altogether. Loss of function of Kmg, Dany, or Simj also resulted in increased expression from Aly-bound, Aly-dependent promoters that normally produced only low levels of transcripts in wild type. ChIP-seq for Kmg showed enrichment at these low expressed promoters, suggesting that Kmg may act locally to repress productive transcription from weak Aly-dependent promoters. This is consistent with the finding from Stampfel *et al*. 2015 that Kmg (CG5204) acted as a powerful repressor when artificially tethered next to a core promoter (Stampfel *et al*. 2015). ChIP-seq also revealed Kmg bound at cryptic promoters when function of Mi-2 or Dany was knocked down in spermatocytes. This observation suggests that Kmg may act transiently at cryptic promoters in wild type testes, and can only be seen enriched at cryptic promoters when its binding partners are not functional and the complex fails to repress and close chromatin. Strikingly, Kmg, Dany, Mi-2, and Simj did not appear to repress productive transcription from Aly-dependent promoters that were highly active in wild type spermatocytes. ChIP-seq from wild type testes revealed that Dany and Mi-2 colocalized with Kmg along the gene bodies rather than at the promoter of highly expressed Aly-dependent genes. When function of Mi-2 was knocked down in spermatocytes, localization of Kmg along the gene bodies of highly expressed Aly-dependent genes dramatically decreased, and transcript levels were downregulated. Based on recent findings by Ullah *et al*. (2022) that binding of Mi-2 to RNA antagonizes its ability to bind and act on chromatin, we discuss a model where binding of Mi-2 to nascent transcripts at highly expressed genes pulls associated Kmg/Dany and NuRD complex components away from the promoter, allowing continued high expression (Ullah *et al*. 2022). In this way the Kmg/Dany/Mi-2 cell-type specific surveillance machinery may provide a mechanism to enhance differences in output between highly expressed vs. weak or cryptic promoters, acted upon in the same cell type by the tMAC complex.

## Results

### Kmg keeps thousands of cryptic promoters repressed genome-wide

Full transcriptome analysis revealed that Kmg is required in spermatocytes to keep transcription repressed from thousands of cryptic promoters across the genome. To gain a full picture of the effects of loss of Kmg on transcription in spermatocytes, we utilized PacBio Long-Read RNA sequencing (Iso-seq), coupled with Short-Read RNA-seq, CAGE and 3’ end seq, to generate a full and accurate transcriptome from wild type testes and testes lacking Kmg function (Figure S1). Kmg loss of function testes were from flies expressing a UAS-RNAi against Kmg, with spermatocyte-specific knockdown induced by the *bamGal4* driver. Control testes were from flies carrying the *bamGal4* driver, without the RNAi construct and grown under the same conditions as the knockdown. Because the function of Kmg seemed to be to control which promoters were expressed in spermatocytes, promoter expression was quantified by summing the expression of every transcript starting from a given promoter region, as defined by CAGE cluster (Figure S1). In Kmg knockdown testes, 5490 promoters were upregulated more than 4-fold compared to control testes (Figure 1B, light pink). Applying more stringent criteria, 2065 promoters were upregulated more than 16-fold and expressed among the top 75% of promoters in the Kmg knockdown (higher than quantile 0.25). Of these, only 84 promoters overlapped with transcription start sites (TSS) previously annotated in the *Drosophila* genome. Removing these from the list resulted in a total of 1981 stringently defined cryptic promoters that fired in Kmg knockdown testes (Figure 1B, dark pink). These 1981 cryptic promoters mapped all over the major autosomes, except to pericentromeric regions. Cryptic promoters were less represented on the X chromosome, and mostly absent from the Y and the 4^th^ chromosome (Figure S2).

### Expression from cryptic promoters may profoundly alter the male germ cell proteome

The over 3000 polyA+ transcript isoforms expressed in Kmg knockdown testes from the 1981 stringently defined cryptic promoters very often shared downstream exons and 3’ ends with annotated genes, a feature that may contribute to proper termination, polyadenylation and/or stability of the cryptic transcripts. Most (86%) of the cryptic transcripts were predicted to encode polypeptides and of those, 90% shared exons with annotated genes (Figure 1C). In 25% of the cases where cryptic transcripts shared downstream exons with annotated genes, the host genes were not expressed in wild type testes (360 genes) (Figure 1D). For example, although *CG6296* mRNA was not detected in control testes, in Kmg knockdown testes, two different cryptic promoters downstream of the annotated promoter fired, driving expression of two different cryptic transcripts (Figure 1E). Both were predicted to encode polypeptides in frame with the host gene, resulting in N-terminally truncated isoforms of the CG6296 protein. RNA FISH using probes for exons common to the normal and the cryptic transcript isoforms confirmed that *CG6296* mRNA was not detected in control testes, but was abundantly expressed specifically in spermatocytes when Kmg was knocked down (Figure 1F,G). For the majority (75%) of cryptic transcripts that shared exons with annotated genes, the host gene was expressed in wild type testes (1008 genes) (Figure 1D). Of the 14% of cryptic transcripts (461) not predicted to encode a polypeptide, only 22% matched with annotated genes, while the majority (78%) were scored as novel (Figure 1C). Of these, half mapped to the antisense strand of annotated genes, and the other half mapped to intergenic regions.

Of the cryptic transcripts that spliced or fused into exons of annotated genes, analysis of their protein coding potential compared to their host gene, predicted 1416 cryptic protein isoforms (Figure 1H). Most of these (97.6%) were in frame with the host protein coding sequence, thus related to proteins encoded by 1136 annotated genes. In most cases (77%) the cryptic proteins were truncated compared to the proteins encoded by the host locus. Analysis by Western blot confirmed several cases of expression of in frame truncated proteins from cryptic transcripts expressed in Kmg knockdown testes. In control testes, *Taf1* was expressed from two different promoters, one of which was not previously annotated in the *Drosophila* genome (Figure 1I, grey transcripts). In Kmg knockdown testes, firing of a cryptic promoter located much further downstream resulted in expression of two much shorter transcripts that spliced into downstream exons from the *Taf1* locus (Figure 1I, pink transcripts). These cryptic transcripts contained an AUG near the 5’ end in frame with the Taf1 protein sequence, predicting expression of smaller proteins of 102KD and 98KD containing the C-terminal domains of Taf1. Probing Western blots of control testis extracts with anti-Taf1 detected the expected large band of around 250KD. Additionally, a smaller band the size of the predicted truncated proteins was detected in Kmg knockdown testis extracts (Figure 1J). Similar analysis with antibodies against Chromator (Chro) and Half pint (Hfp) also detected truncated proteins of the predicted size expressed in Kmg knockdown testes, but not in control testes (Figure 1J; Figure S3).

### Kmg interacts with Dany, Mi-2 and other NuRD complex components to keep cryptic promoters repressed

Immunoprecipitation followed by Mass Spectrometry (IP-MS) identified proteins that work with Kmg to block expression from cryptic promoters. Immunoprecipitation of Kmg from *Drosophila* testes recovered the large chromatin remodeler Mi-2 (homolog of mammalian Chd3/4/5) as expected from previous work (Kim *et al*. 2017). Anti-Kmg IP-MS also identified other components of the Nucleosome Remodeling and Deacetylase (NuRD) complex, including Simj (p66/Gatad2a/b), CG15142 (a testes specific paralog of CDK2AP1/DOC-1), and to a lesser extent HDAC1 (Rpd3) (Figure 2A, complete list in Table S1). In addition, the anti-Kmg IP-MS also recovered peptides from Dany, implicated in regulation of gene expression in *Drosophila* spermatocytes (Trost *et al*. 2016). Dany, like Kmg, is expressed only in spermatocytes, with onset of Dany protein expression starting very early in the spermatocyte stage (Figure 2B). Dany and Kmg proteins colocalized to spermatocyte nuclei along with Mi-2 protein (Figure 2B), which is more generally expressed. Co-Immunoprecipitation confirmed that Kmg binds Dany and Mi-2. Immunoprecipitation of Mi-2-GFP from testes expressing a Dany allele tagged with V5 at the endogenous locus recovered both Kmg and Dany proteins (Figure 2C).

**Figure 2:**
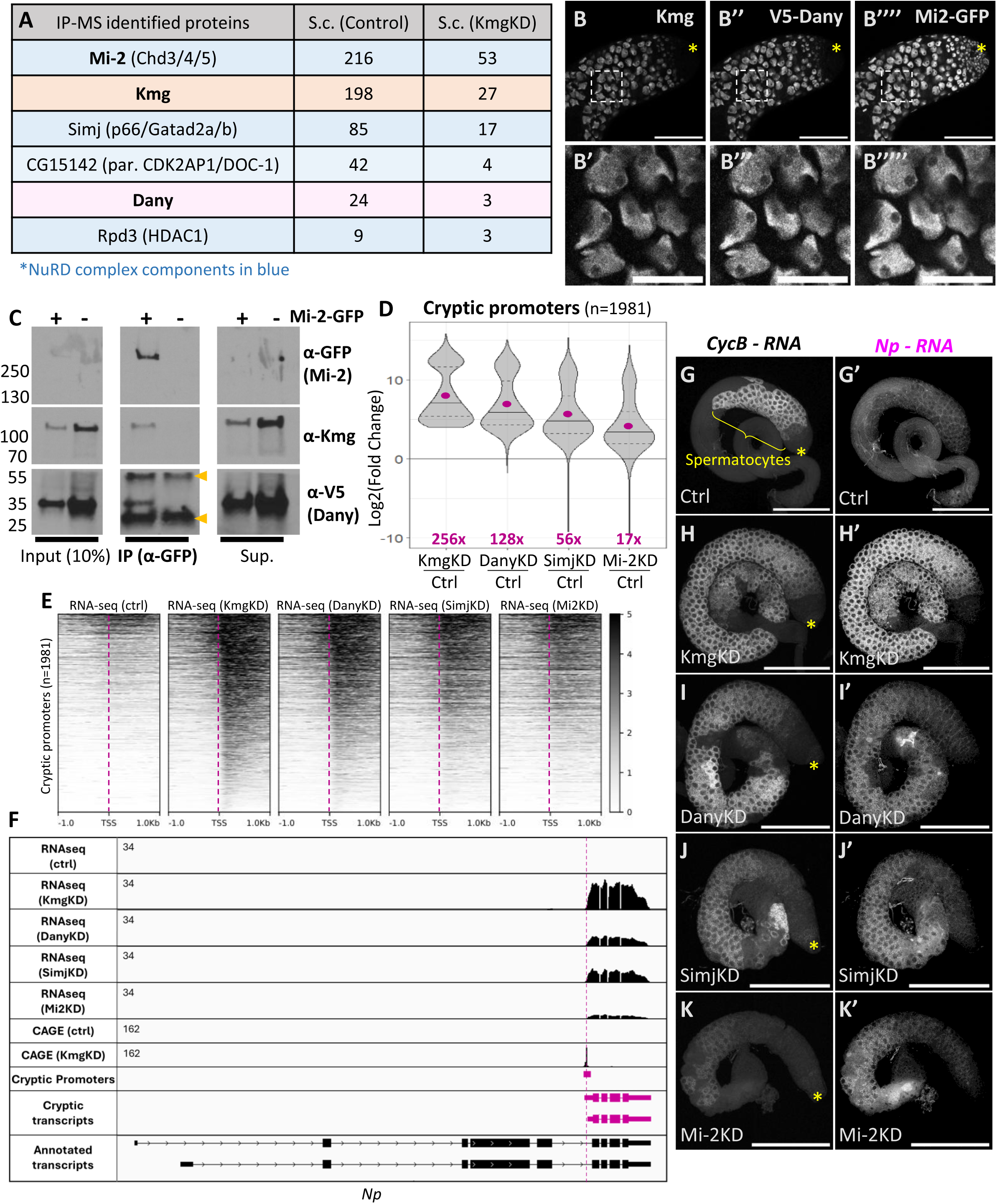
Kmg interacts with Dany and Mi-2 (NuRD) to repress cryptic promoters. (A) Spectral counts by protein from immunoprecipitation of Kmg followed by Mass Spectrometry (IP-MS), from control or Kmg KD testes. Blue: NuRD complex. (B) Immunofluorescence images showing apical tip of a testis expressing V5-Dany and Mi-2-GFP, stained with anti-Kmg, anti-V5, and anti-GFP. Asterisk: testis apical tip. Scale bars, 100 µm. (C) Immunoprecipitation (IP) with anti GFP from testes expressing V5-Dany and either Mi-2-GFP (+), or not (-), followed by PAGE and Western blots probed with anti-GFP, anti-Kmg and anti-V5. Yellow arrow heads point to bands corresponding to the immunoglobulin heavy (around 50 kDa) and light (around 25 kDa) chains from anti-GFP, which was cross-linked to the IP beads. (D) Violin plot of log2 transformed fold change for the 1981 cryptic promoters in Kmg KD, Dany KD, Simj KD and Mi-2 KD, compared to control testes. Pink dot: mean value (not log2 transformed, indicated at the bottom). Excluding 20 out of scale promoters. (E) Heatmap plot of normalized RNA-seq signal for control, Kmg, Dany, Simj and Mi-2 knockdown testes, showing +/-1Kb centered at the TSS of the highest expressed transcript starting at each cryptic promoter, aligned 5’ to 3’ relative to each transcript. Loci sorted by RNA-seq signal in Kmg knockdown, highest on top. (F) RNA-seq and CAGE profiles for the *Np* gene. Pink: cryptic promoters and transcripts (G-K) HCR-FISH using probes against *CycB* or cryptic *Np* RNA, in control (G,G’), Kmg (H,H’), Dany (I,I’), Simj (J,J’) and Mi-2 (K,K’) knockdown testes.

Function of Dany and the NuRD complex components Simj and Mi-2, was required to block expression from the same cryptic promoters that require Kmg. Analysis of promoter expression in testes expressing UAS-RNAi against either Dany, Simj or Mi-2 in spermatocytes under control of *bamGal4*, revealed that all three proteins were needed to repress the same cryptic promoters (Figure 2D, S4). These cryptic promoters were highly upregulated in Dany knockdown testes (128x), as in Kmg knockdown (256x). Expression of cryptic promoters was also elevated in Simj (56x) or Mi-2 (17x) knockdown testes although to a lesser extent. Indeed the genomic coordinates of cryptic promoters, defined using Kmg knockdown samples, also marked initiation of cryptic transcription in Dany, Simj and Mi-2 knockdown testes (Figure 2E). For example, the expression of the *Notopleural* (*Np*) gene was not detected by RNA-seq in control testes. However, testes in which function of Kmg, Dany, Simj or Mi-2 was knocked down showed expression from the same cryptic promoter, considerably downstream of the annotated promoter for *Np* (Figure 2F). RNA FISH with probes recognizing the cryptic transcripts of *Np* detected expression in spermatocytes of the different loss of function testes, but not in control testes (Figure 2G-K).

### In Kmg knockdown testes, cryptic promoters are bound by Aly and are Aly-dependent

In control testes, expression from cryptic promoters was either not detected, or was at extremely low levels (Figure 1B). Analysis by ChIP-seq of testes from flies containing a copy of transgene with epitope tagged Aly (Aly-HA), showed that in control testes very little Aly protein (a component of the tMAC activator complex) was detected at cryptic promoters (Figure 3A,B). ATAC-seq in control samples enriched for mid-stage spermatocytes (ctrl^72hrs^), and CAGE in control testes, revealed that the chromatin at these loci was largely not accessible and little transcription initiation occurred at the cryptic promoter sites (Figure 3A,C,D,E). In Kmg knockdown testes, in contrast, ChIP-seq for Aly-HA revealed strong enrichment at cryptic promoter sites, which were accessible to ATAC-seq and had robust transcription initiation revealed by CAGE (Figure 3A-E). Plots of two genomic regions with cryptic promoters illustrated the increase in Aly ChIP-seq signal at cryptic promoter sites in Kmg knockdown testes. At the *CG10628* locus, loss of function of Kmg resulted in firing of a cryptic promoter upstream of the normally utilized promoter (Figure 3F). A cryptic promoter at the *krotzkopf verkehrt* (*kkv*) locus drove the expression of a cryptic transcript in the antisense direction of the host gene in Kmg knockdown testes (Figure 3G). In both cases, binding of Aly at the cryptic promoter site increased, new CAGE signal appeared, and the cryptic promoter became more accessible to ATAC-seq in Kmg knockdown testes.

**Figure 3:**
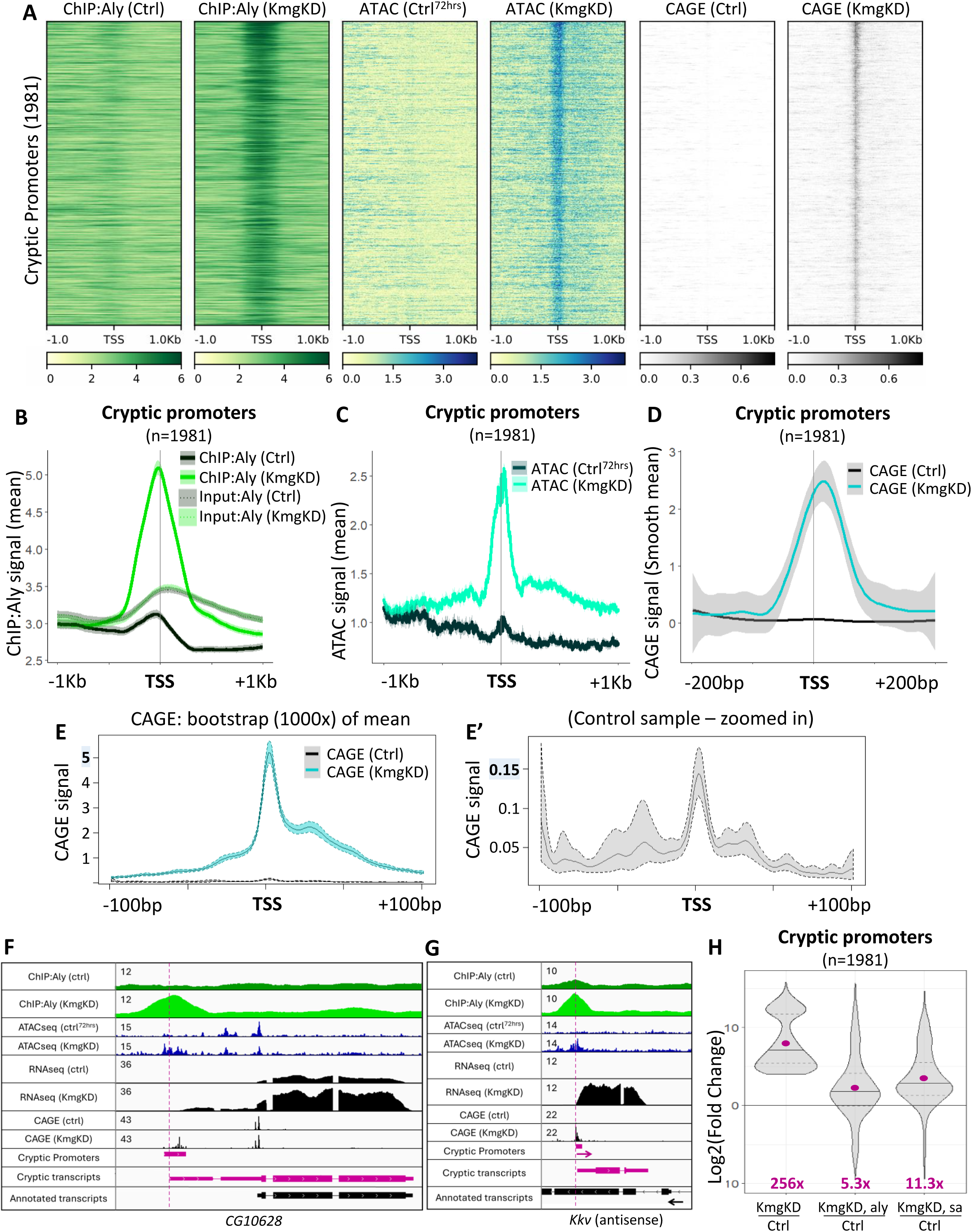
In absence of Kmg, cryptic promoters are enriched by Aly and depend on Aly for expression. (A) Heatmap plots of ChIP-seq for Aly-HA, ATAC-seq and CAGE in control and Kmg KD testes, showing +/-1Kb centered at the TSS of the highest expressed transcript starting at each cryptic promoter, aligned 5’ to 3’ relative to each transcript. Loci sorted by level of promoter expression, highest on top. (B-D) Mean profiles of (B) Aly-HA ChIP-seq (solid lines) and corresponding input (dotted lines), (C) ATAC-seq and (D) CAGE signals centered around the same regions as in (A). (D) CAGE mean signal was smoothed by method = “gam”. (E) Bootstrapping analysis of CAGE data, with 1000x iterations. (E’) Control sample with zoomed in scale on Y axis. (F-G) Aly-HA ChIP-seq, ATAC-seq, RNA-seq and CAGE profiles for the *CG10628* (F) and *Kkv* (G) genes. Pink: cryptic promoters and transcripts. (H) Violin plot of log2 transformed fold change compared to control, for the 1981 cryptic promoters in Kmg knockdown testes, Kmg knockdown in aly mutant testes, Kmg knockdown in sa mutant testes. Pink dot: fold change mean value (not log2 transformed, indicated at the bottom). Excluding 14 out of scale promoters on H.

Analysis of promoter expression showed that transcription from cryptic promoters depended on function of Aly and the testis TAF Sa for maximal transcript production. Expression from cryptic promoters was considerably lower in testes lacking both Kmg and Aly than for loss of Kmg function alone (Figure 3H). Expression from cryptic promoters was also lower in testes lacking both Kmg and Sa (Figure 3H), a testis-specific TAF (tTAF) known to boost expression of tMAC dependent genes in wild type testes (M. Hiller *et al*. 2004). Together these data showed that, in Kmg knockdown testes, cryptic promoters were bound by Aly and their expression was dependent on Aly and Sa function for full expression.

### Kmg is enriched at cryptic promoters in Mi-2 and Dany knockdown testes

Although Kmg did not accumulate at cryptic promoter sites in control testes, Kmg protein became enriched at cryptic promoters in spermatocytes lacking function of Dany or Mi-2 (Figure 4A,B). As discussed later, we suspect that Kmg acts transiently to repress cryptic promoters in wild type testes, but is ineffectual and stays bound at promoters in the absence of function of its binding partners.

**Figure 4:**
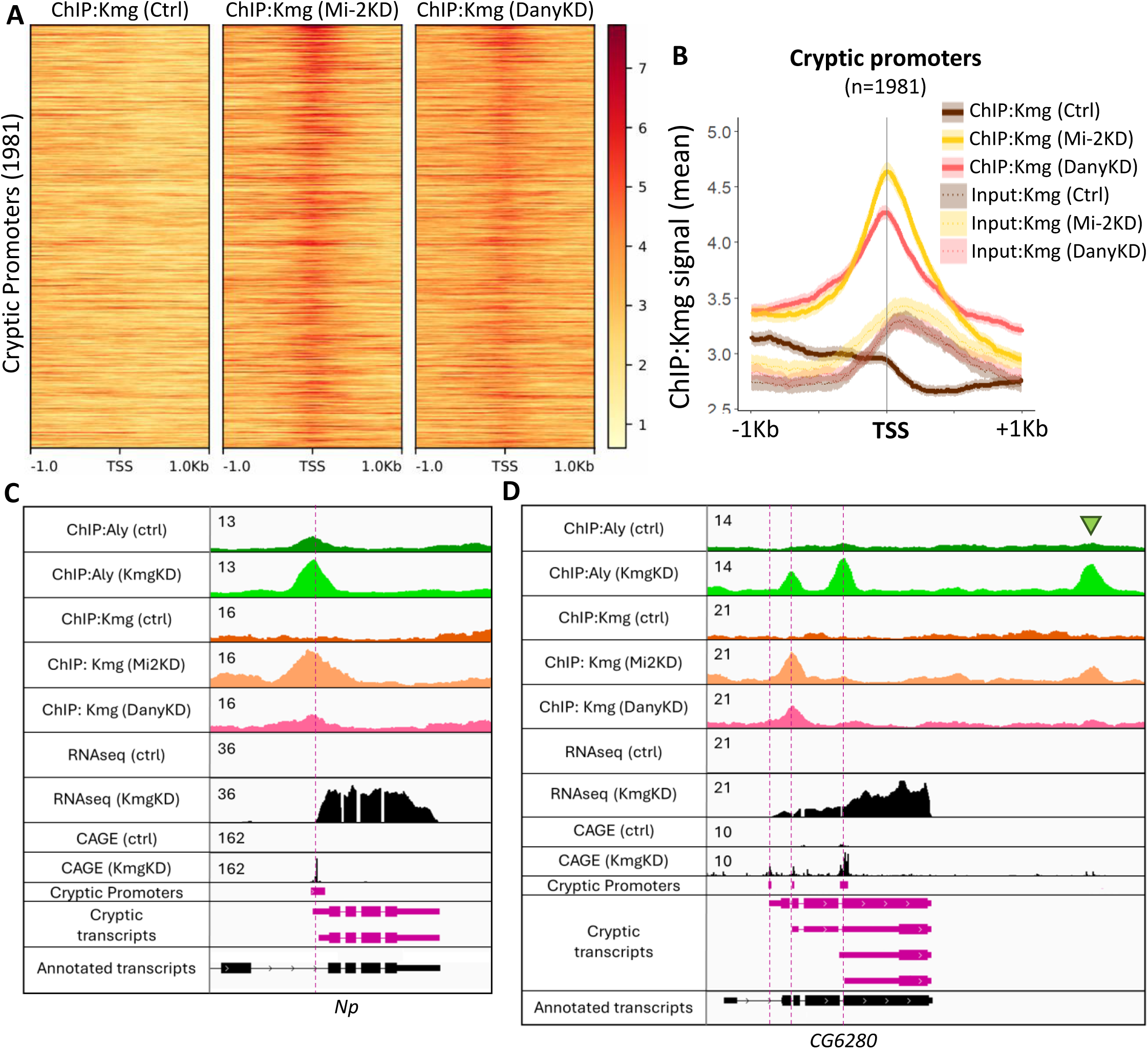
Kmg is enriched at cryptic promoters in Mi-2 and Dany knockdown. (A) Heatmap plots of ChIP-seq for Kmg in control, Mi-2 and Dany knockdown testes, showing +/-1Kb centered at the TSS of the highest expressed transcript starting at each cryptic promoter, aligned 5’ to 3’ relative to each transcript. Loci sorted by level of promoter expression, highest on top. (B) Mean profiles of Kmg ChIP-seq (solid lines) and corresponding input (dotted lines) signals centered around the same TSS regions as in (A). (C-D) Aly-HA ChIP-seq, Kmg ChIP-seq, RNA-seq and CAGE profiles for (C) the *Np* and (D) the *CG6280* genes, in control, Kmg, Dany or Mi-2 knockdown testes.

is recruited and stays in place in the absence of function of its binding partners.

Plots of ChIP-seq, RNA-seq and CAGE data at two example genes with cryptic promoters showed no accumulation of ChIP-seq signal for Kmg at the cryptic promoters in control testes, but substantial enrichment of Kmg when function of either Mi-2 or Dany was knocked down in spermatocytes (Figure 4C,D). Loss of function of Kmg resulted in increased signal from ChIP-seq for the tMAC component Aly at these loci (Figure 4C,D), as shown for other cryptic promoters in Figure 3.

Kmg prevented Aly from binding thousands of loci genome wide that did not overlap with productive promoters. In addition to the novel Aly peaks that appeared at cryptic promoters in Kmg loss of function testes, 6,712 Aly peaks (peak q value < 10^−10^) that did not overlap with promoters were called in Kmg knockdown testes, but not in control samples (Figure S5A) (see Figure 4D green arrow for an example). Chromatin dynamics at these non-productive Aly peaks closely resembled that at cryptic promoters. While these sites showed little enrichment by ChIP-seq for Aly and little ATAC-seq signal in control testes, they became bound by Aly and accessible to ATAC-seq in Kmg knockdown testes (Figure S5A,B). Although production of PolyA+ transcripts was not detected from these loci in Kmg knockdown testes, a very low enrichment for CAGE signal was observed. (Figure S5A,C). It is possible that transcription weakly initiated but was not successful at these open chromatin regions because the DNA lacked sequences required for robust recruitment of Pol II, Mediator, testes TAFs, or other effectors needed for transcription elongation (C. Lu and Fuller 2015; M. A. Hiller *et al*. 2001; M. Hiller *et al*. 2004). Alternatively, transcripts initiating from these loci could be prematurely terminated, fail to be polyadenylated, or be rapidly degraded.

The Aly peaks that appeared at these non-promoter sites upon loss of function of Kmg marked regions that became enriched for Kmg in testes lacking function or Mi-2 or Dany (Figure S5A,D). Together, these results showed that loss of function of Kmg in spermatocytes resulted in increased binding of the tMAC component Aly at many sites scattered across the genome. Some of these new Aly-bound sites produced polyA+ transcripts from normally silent (cryptic) promoters, while many other sites with novel Aly peaks did not substantially produce transcripts. Both the productive and the non-productive novel Aly peaks, however, were enriched for Kmg protein in spermatocytes lacking Mi-2 or Dany.

### Kmg is enriched at loci bound by Aly in wild type testes

In wild type testes, the tMAC component Aly also bound to thousands of sites across the genome, many of which did not initiate appreciable levels of PolyA+ transcripts. ChIP-seq for Aly identified 14,400 peaks strongly enriched by Aly in control testes (peak q value < 10^−10^), of which 4,786 Aly peaks (33%) overlapped with promoters expressed in control testes. For 1,952 of these, expression from the promoters was strongly Aly-dependent (downregulated more than 16-fold in *aly* mutants compared to control) (Figure 5A, top; Figure S6). Surprisingly, Aly was enriched at many more sites on the genome than the promoters it regulates. 1079 Aly peaks mapped to promoters that were expressed in testes but did not require Aly for expression (Figure 5A, middle, Figure S6). In addition, Aly was robustly enriched at 9,585 loci (67% of the total Aly peaks) that did not overlap with promoters detected as expressed in testes (neither in control, nor in Kmg knockdown testes) (Figure 5A, bottom). Similar to the novel non-transcribing Aly-bound loci observed in Kmg knockdown testes (Figure S5A,B), these Aly peaks observed in wild type testes may lack important promoter information for transcription initiation or elongation, or produce unstable RNAs. Comparing ATAC-seq data from *aly* mutant testes to testes samples enriched for mid-stage spermatocytes (ctrl^72hrs^) showed that Aly was needed to open chromatin at both the Aly-dependent promoters (Figure 5A,D) and the thousands of loci that had Aly peaks but lacked productive transcription (Figure 5A,D,F). In contrast, function of Aly was not required for chromatin to be open at Aly-bound but Aly-independent promoters, which expressed similar levels of transcripts in *aly* mutant testes as in wild type (Aly-independent promoters, Figure 5A,E). As expected, CAGE signal was detected at loci where productive elongation was detected (Figure 5A,G,H). Aly peaks that did not overlap with productive promoters had very low detectable CAGE signal (Figure 5A,I).

**Figure 5:**
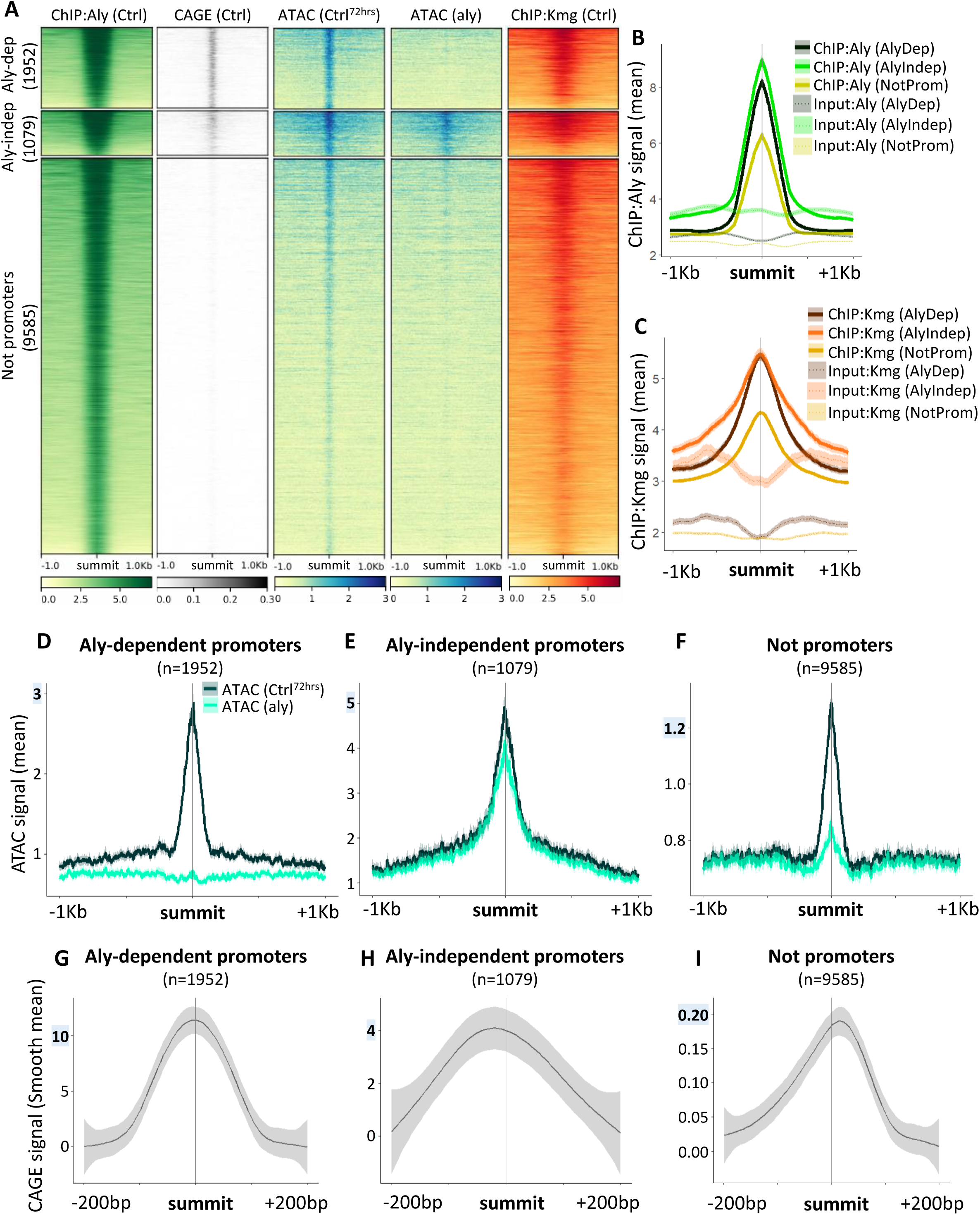
Kmg is enriched at loci bound by Aly in control testes. (A) Heatmap plots of ChIP-seq for Aly-HA, ChIP-seq for Kmg and CAGE in control testes, ATAC-seq in control testes enriched for mid-stage spermatocytes (Ctrl^72hrs^) and aly mutant testes. Plots are centered around the summit of each Aly peak called in control testes (peak q value < 10^−10^), +/-1Kb. Loci sorted by the level of Aly-HA ChIP-seq signal in control testes. Aly peaks were divided into 3 groups: Aly-dependent promoters (1952 peaks), Aly-independent promoters (1079 peaks), Not on promoters (9585 peaks). See figure S6 for cutoffs, and methods for promoter assignment. (B-C) Mean profiles of ChIP-seq results for (B) Aly-HA and (C) Kmg (solid lines) and corresponding input (dotted lines) centered on the same regions as in (A). (D-F) Mean profiles of ATAC-seq signals centered as in (A). (G-I) Mean smoothed by method = “gam”, profiles of CAGE signals centered as in (A), +/-200bp. (D-I) Note: Y axis scales differ between the Aly peak groups.

Kmg colocalized with peaks of Aly throughout the genome in wild type testes, whether or not those Aly peaks overlapped with promoters. ChIP-seq for Kmg protein showed enrichment at most of the loci robustly bound by Aly in control testes (Figure 5A,C), with the level of Kmg ChIP-seq signal seeming to correlate with the level of Aly ChIP-seq signal (Figure 5A,B,C).

### Kmg, Dany and Mi-2 localize along the transcribed region downstream of highly expressed Aly-dependent promoters

While ChIP-seq for Kmg enriched for sequences near the transcription start site of most Aly-dependent promoters, the results were different for very highly expressed Aly-dependent promoters, where Kmg co-localized with Dany and Mi-2 along the gene body (Figure 6A). The set of promoters that bound Aly (ChIP-seq) and required Aly function to be accessible (ATAC-Seq) and initiate transcription (CAGE signal) (1952 Aly-dependent, in Figure 5A, top), were transcribed at various levels, from very high to low (RNA-Seq signal) (Figure 6A). For example, promoters for the *CG4836* and *S-Lap7* genes were bound by Aly and were among the highest expressed Aly-dependent promoters transcribed in spermatocytes. Analysis of ChIP-seq at these loci showed a peak of Aly at the promoter regions, located just upstream of the TSS, while ChIP-seq for the Kmg, Dany or Mi-2 proteins enriched for sequences along the gene body (Figure 6B). These binding profiles were general features of the top 5% highest expressed Aly-dependent promoters (Figure 6C).

**Figure 6:**
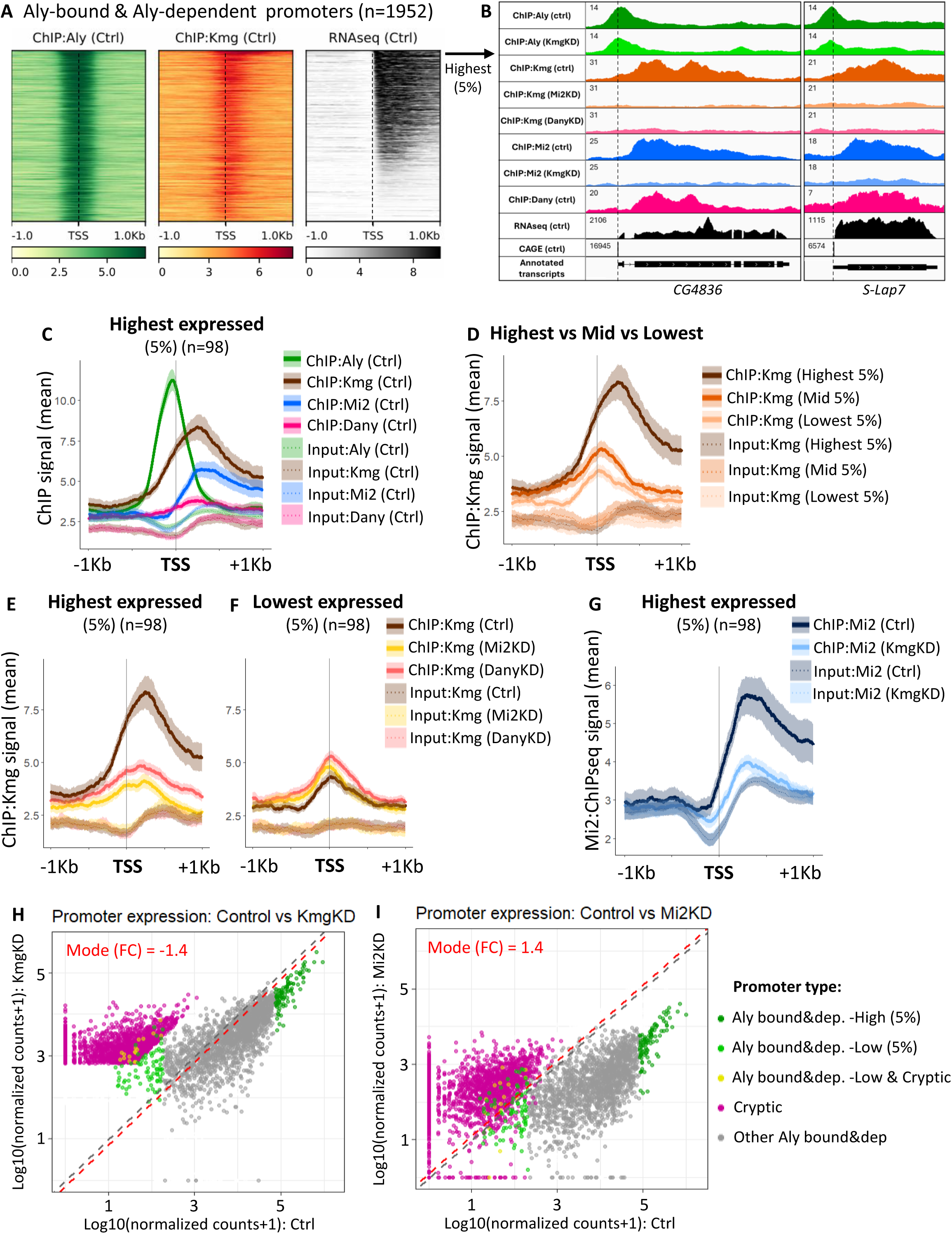
Location of the Kmg complex at Aly-bound, Aly-dependent promoters depends on expression level. (A) Heatmap plots of ChIP-seq for Aly-HA and Kmg, and RNA-seq in control testes, showing +/-1Kb centered at the TSS of the highest expressed transcript starting at each Aly-dependent promoter (defined in Figure 5), aligned 5’ to 3’ relative to each transcript. Loci sorted by level of promoter expression, highest on top. (B) ChIP-seq data for Aly-HA, Kmg, Dany and Mi-2, RNA-seq and CAGE profiles for *CG4836* and *S-Lap7* loci. (C) Mean profiles of ChIP-seq results for Aly-HA, Kmg, Dany and Mi-2 (solid lines) and corresponding input (dotted lines) signals for the top 5% highest expressed promoters, centered on the TSS and aligned 5’ to 3’. (D) Mean profiles of Kmg ChIP-seq (solid lines) and corresponding input (dotted lines) results comparing the top 5% highest, Mid 5% and 5% lowest expressed promoters, centered on the TSS. (E-F) Mean profiles of Kmg ChIP-seq (solid lines) and corresponding input (dotted lines) results, in control, Mi-2 or Dany knockdown testes, for the (E) top 5% highest or (F) 5% lowest expressed promoters, centered on the TSS. (G) Mean profiles of Mi-2 ChIP-seq (solid lines) and corresponding input (dotted lines) results, in control or Kmg knockdown testes, for the top 5% highest expressed promoters. (H-I) Expression profile for Aly-bound, Aly-dependent promoters (dark green: top 5% highest, light green: 5% lowest, grey: others), cryptic promoters (pink) and Aly-bound, Aly-dependent promoters that are also classified as cryptic promoters (yellow). Axes: log10 transformation of read counts normalized by DESeq2 in testes from control [X-axis] compared to (H) Kmg knockdown [Y axis], or (I) Mi-2 knockdown [Y axis].

In contrast, for the 5% lowest expressed Aly-bound and Aly-dependent promoters, ChIP-seq for Kmg enriched for sequences near the TSS and not along the gene body in wild type testes. Comparison of Kmg ChIP-seq signal at the 5% most highly expressed, 5% mid expressed and the 5% lowest expressed promoters showed a gradual shift of Kmg distribution, from stretched out along the gene body at highly expressed genes, to a peak near the promoter in lower expressed promoters (Figure 6D).

Strikingly, binding of Kmg along the gene body downstream of highly expressed Aly-dependent promoters required function of the Kmg-interacting proteins Dany and Mi-2. When expression of Dany or Mi-2 was knocked down in spermatocytes, the Kmg ChIP-seq signal was much reduced along the gene bodies of *CG4836* and *S-Lap7* (Figure 6B). In general, at the 5% most highly expressed Aly-bound and Aly-dependent promoters, knockdown of Dany or Mi-2 resulted in loss of Kmg ChIP-seq signal from along gene bodies, and some enrichment of Kmg near promoters (Figure 6E). At the lowest expressed Aly-dependent promoters, however, ChIP-seq for Kmg enriched for sequences at the TSS in control testes, in a pattern that remained largely the same upon loss of function of either Dany or Mi-2 (Figure 6F).

### Mi-2 loss of function results in downregulation of highly expressed Aly-dependent promoters

Although Kmg, Dany and Mi-2 colocalized at gene bodies downstream of highly expressed Aly-dependent promoters, loss of function of these proteins had different effects on promoter expression. Highly expressed Aly-bound and Aly-dependent promoters were still expressed at very high levels in Kmg knockdown or Dany knockdown testes compared to control testes (dark green dots) (Figure 6H, Figure S7B,C). Expression from the promoters appeared slightly lower than in control testes, in part due to the higher complexity of the Kmg and Dany loss of function RNA-seq libraries compared to control (Figure S8). Similarly, ChIP-seq for Aly showed only slightly less enrichment at these promoters in Kmg knockdown compared to control testes (Figure S7E). In contrast, loss of function of Mi-2 in spermatocytes resulted in dramatic downregulation of the highly expressed promoters (Figure 6I, Figure S7C). Furthermore, ChIP-seq for Mi-2 did not enrich along the gene body of highly expressed loci in Kmg knockdown testes (Figure 6B,G), suggesting Kmg function was necessary for recruitment of Mi-2 to chromatin at these sites. However, despite the lack of local Mi-2 along the gene bodies downstream of highly expressed promoters in Kmg knockdown testes, expression from the promoters remained high in Kmg knockdown. We speculate that Mi-2 may be required at highly expressed loci to prevent repression by Kmg (see discussion).

### Kmg acts to dampen transcription of lowly expressed Aly-dependent promoters

In contrast to highly expressed promoters, function of Kmg and Dany in spermatocytes was required to keep expression from weakly expressed Aly-dependent promoters low. As the level of expression from Aly-bound and Aly-dependent promoters decreased in control testes (Figure 6H, X axis), their level of expression in Kmg knockdown testes increased compared to control testes (Figure 6H, Y axis). A similar gradual shift was observed for Dany knockdown testes (Figure S7A). The Aly-bound and Aly-dependent promoters expressed at low levels in control testes (light green dots) were strongly upregulated in Kmg knockdown and Dany knockdown compared to control testes (Figure 6H, S7A,D). Kmg and Dany appeared to have similar but more extreme effects at cryptic promoters (pink dots), which were even less expressed in control testes, and strongly upregulated upon loss of function of Kmg or Dany (Figure 6H, S7A). Indeed, some Aly-bound, Aly-dependent promoters expressed at very low levels in control testes became so highly upregulated in Kmg knockdown testes that they reached the threshold to be classified as cryptic promoters (Figure 6H, yellow dots). For the Aly-bound, Aly-dependent promoters expressed at low levels in wild type, as well as the cryptic promoters, chromatin accessibility and expression from the promoter required function of Aly (Figure 3, Figure 5).

## Discussion

Our results show that the transcription program that drives differentiation in the *Drosophila* male germ line adult stem cell lineage, set up by the master regulator of spermatocyte differentiation tMAC, is restrained by a cell-type specific mechanism that downregulates weak promoters while allowing robust transcription from highly expressed, tMAC dependent genes. Based on ChIP-seq for its subunit Aly, the spermatocyte-specific tMAC complex binds thousands of sites in the genome, where in most cases Aly function is required for chromatin to become accessible from a characteristically closed state in spermatogonia. Robust transcription initiates from a subset of these Aly-bound loci, in many cases triggering genes only expressed in spermatocytes, and in others firing alternative, spermatocyte-specific promoters in genes already being expressed in previous developmental stages or other cell types (this study, Lu *et al*., 2020).

We found that Kmg and Dany, two proteins expressed at the onset of spermatocyte development, interact with the nucleosome remodeler Mi-2, and possibly other components of the NuRD complex, to dampen productive transcription from normally low expressed Aly-dependent promoters (Figure 7A). ChIP-seq for Kmg enriched for Aly-bound, Aly-dependent, low expressed promoters, and transcription from these promoters was upregulated upon loss of function of Kmg, Dany, or the NuRD component Simj. This, together with the previous finding that Kmg scored as one of the most powerful repressors when artificially tethered to different core promoters or enhancers (Stampfel *et al.,* 2015), suggests that Kmg could be acting as a direct repressor of transcription at Aly-bound promoters in spermatocytes.

**Figure 7:**
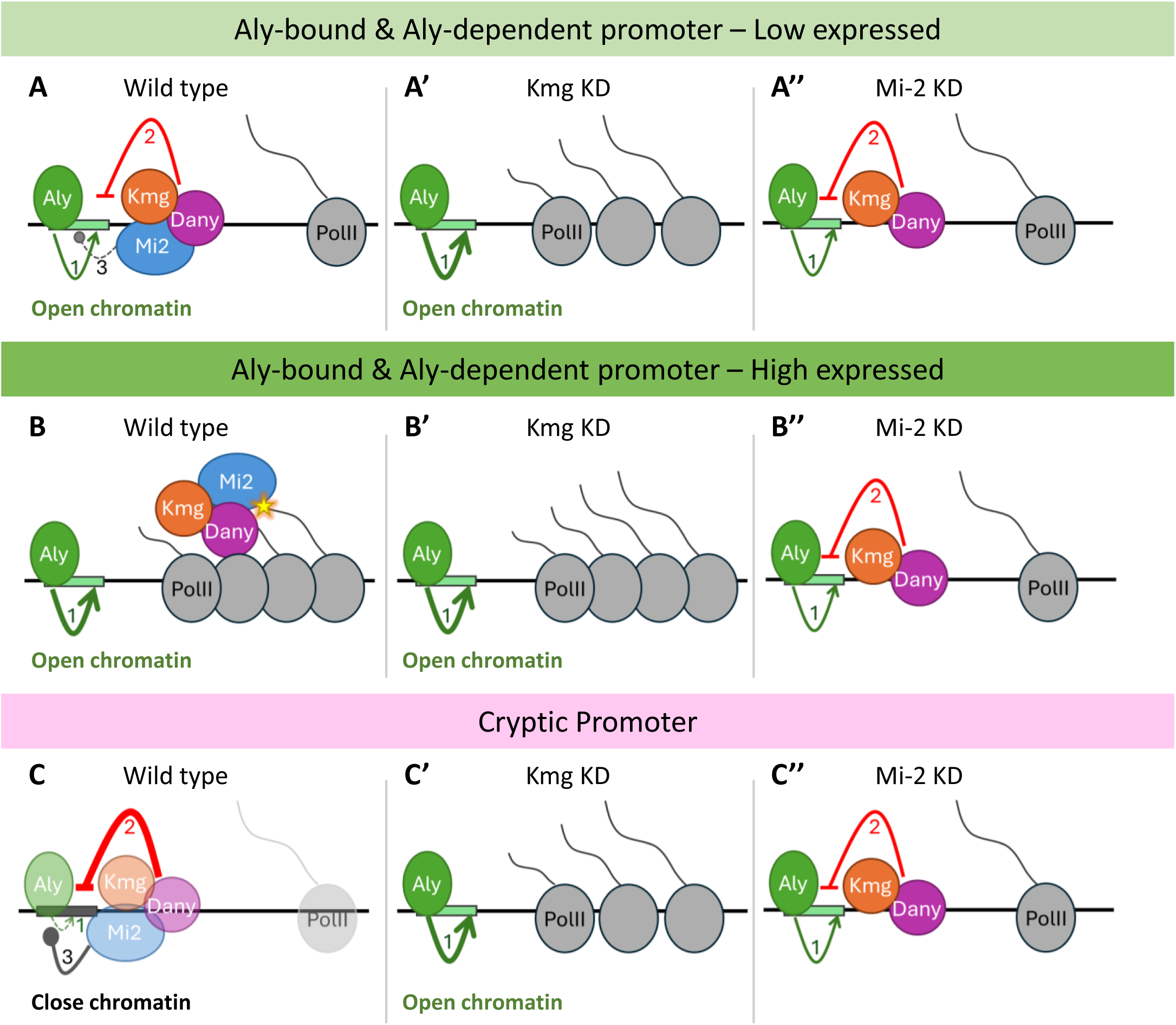
Proposed model. (A,A’,A’’) Diagram of an Aly-bound, Aly-dependent promoter expressed at low levels in wild type. (A) In wild type testes, Aly (tMAC) binds to the promoter and opens chromatin allowing expression (1). The Kmg/Dany complex at the promoter site dampens transcription, keeping expression low (2). We propose that Mi-2, a chromatin remodeler, is not successful in closing chromatin at these sites (3). (A’) In Kmg knockdown testes, the Kmg complex is not recruited, or not functional, chromatin is open and the promoter upregulated (1). (A’’) We propose that in Mi-2 knockdown testes, chromatin does not close, and expression level of the promoter stays low, as Kmg/Dany can still repress transcription (2). (B,B’,B’’) Diagram of an Aly-bound, Aly-dependent promoter expressed at very high levels in wild type. (B) We speculate that in wild type testes, Mi-2 associates with nascent RNAs transcribed from the very active promoters. This (yellow star) antagonizes Mi-2 binding to chromatin and allows Kmg/Dany to be pulled away from the promoter, which is then protected from repression and highly active (1). (B’) In Kmg knockdown testes, the Kmg complex is not recruited, or not functional, and the promoter is expressed at high levels (1). (B’’) We propose that in Mi-2 knockdown testes, Kmg/Dany are not pulled away from the promoter where it can dampen transcription (2). (C,C’,C’’) Diagram of a cryptic promoter. (C) We speculate that in wild type testes, Aly initially binds to chromatin (1), the Kmg/Dany complex represses transcription (2) and Mi-2 closes chromatin (3) evicting Aly. The binding of these players at cryptic promoters is thus transient. (C’) In Kmg knockdown testes, the surveillance function is lacking, Aly binds and opens chromatin unopposed, and promoters are highly expressed (1). (C’’) We propose that in Mi-2 knockdown testes chromatin does not close (due to loss of the chromatin remodeler), Aly is not evicted, and cryptic promoters are transcribed (1). However, Kmg binding at the promoter site keeps expression at lower levels (2) compared to loss of Kmg.

Action of Kmg, Dany, Simj and Mi-2 is also required to block transcription in spermatocytes from hundreds of cryptic promoters scattered across the major autosomes. These cryptic promoters normally had very little Aly enrichment and remained largely inaccessible to ATAC-seq in wild type spermatocytes, but in absence of Kmg function, Aly bound to the promoter region, a small stretch of local chromatin became accessible, and productive transcription initiated. In spermatocytes lacking function of either Dany or Mi-2, ChIP-seq revealed Kmg enriched at the cryptic promoters. We hypothesize that under normal conditions, both the activating tMAC complex and the Kmg complex may bind transiently at cryptic promoters, with Kmg and Dany effectively repressing transcription and Mi-2 closing chromatin and evicting tMAC (Figure 7C). We speculate that in Dany or Mi-2 loss of function, Kmg is successfully recruited to cryptic promoters, but is not sufficient to repress transcription in the absence of its partner proteins.

Strikingly, Kmg, Dany and Mi-2 were not enriched at Aly-dependent promoters that were highly expressed in wild type, but instead colocalized along the gene bodies of the highly transcribed genes. No components of the RNA Polymerase II complex were detected in our IP-MS for proteins that interact with Kmg. We speculate that the Kmg/Dany/Mi-2 complex may be drawn away from promoters by binding of Mi-2 to nascent RNA, a property of Mi-2 demonstrated by Ullah *et al*. (2022). The authors proposed that binding of Mi-2 to nascent transcripts at highly expressed genes could provide a mechanism to remove the repressive NuRD complex away from promoters, sparing highly expressed genes from repression (Ullah *et al*. 2022). We hypothesize that Mi-2 plays a similar role in the spermatocyte-specific transcription program, clearing the Kmg-Dany complex away from highly active promoters and thus protecting them from repression (Figure 7B). In agreement with this model, when function of Mi-2 was knocked down in spermatocytes, Kmg protein was no longer enriched along the gene body of highly expressed Aly-dependent genes and the promoters were strongly downregulated. Our data suggest that Mi-2 is not required to actively induce expression from the highly expressed, Aly-dependent promoters, because in spermatocytes lacking Kmg function Mi-2 protein was lost from the loci but the promoters were still expressed at very high levels. We suspect that ChIP-seq detected Kmg, Dany, and Mi-2 along very highly expressed gene bodies because the levels of nascent RNA at these loci were so high that the complex could be crosslinked to RNA and adjacent chromatin.

We propose that the same players exist in a dynamic battle at all Aly-bound and Aly-dependent promoters, with the outcome of the battle between chromatin opening and activation by tMAC and repression by the Kmg complex depending on how robustly the promoter is able to drive expression of elongating transcripts. In this framework, the ability of Mi-2 to bind nascent transcripts and so pull the Kmg/Dany repressive complex away from highly active promoters may provide a mechanism to amplify differences in promoter expression levels, suppressing minimal promoters while allowing expression from robust promoters to proceed (Figure 7).

Failure of the Kmg surveillance system results in firing of thousands of cryptic promoters genome wide, generating aberrant transcripts that frequently share downstream exons with known genes, leading to proteomic chaos. Productive elongation from cryptic promoters is predicted to result in expression of potentially hundreds of N-terminally truncated proteins, as well as completely aberrant polypeptides and proteins not normally expressed in spermatocytes. We wonder if similar mechanisms are at play in other key differentiation pathways, and that their failure may contribute to abnormal differentiation or cancer.

## Materials and Methods

### Fly strains and husbandry

All flies were grown on standard molasses media. Fly crosses were kept for 3 days at 25°C. Adults were then flipped to a new bottle, while the eggs and early larvae were shifted to 29°C. For RNAi induced knockdowns (KD), virgins from *bamGal4* (Chen and McKearin 2003) were crossed with males from UAS-*Kmg* RNAi (VDRC-107395), UAS-*Dany* RNAi (VDRC-105571), UAS-*Simj* RNAi (VDRC-100285), or UAS-*Mi-2* RNAi (VDRC-107204). For *aly* mutants, virgins from *aly^5p^*/TM6 were crossed with males from *aly^2^*/TM6, and *aly^2^/aly^5p^* males were dissected (White-Cooper *et al*. 2000). For *sa* mutants, virgins from *sa^1^*/TM6 were crossed with males from *sa^2^*/TM6 and *sa^1^/sa^2^*males were dissected (M. Hiller *et al*. 2004). For Kmg KD and *aly* rescue, virgins from *aly^2^,bamGal4*/TM6 were crossed with males from UAS-*Kmg* RNAi/CyO; *aly^5p^*/TM6 and UAS-*Kmg* RNAi/+; *aly^2^,bamGal4/aly^5p^* males were dissected. For Kmg KD and *sa* rescue, virgins from *sa^1^,bamGal4*/TM6 were crossed with males from UAS-*Kmg* RNAi/CyO; *sa^2^*/TM6 and UAS-*Kmg* RNAi/+; *sa^1^,bamGal4/sa^2^* males were dissected. For Co-IP experiments, flies from the following stocks were dissected: *V5-Dany/V5-Dany*; *Mi-2-GFP*/TM6, or *V5-Dany/V5-Dany*. *Mi-2-GFP* (Flytrap, CA06598, 26). For ChIP-seq of Aly-HA, testes were dissected from flies bearing one copy of the Aly-HA genomic rescue transgene generated by Kim *et al*, 2017.

N-terminal V5 tagged *Dany* (*V5-Dany*) was created by CRISPR-Cas9 mediated V5 sequence insertion at the endogenous *Dany* genomic location. The guide RNA sequence (ACTCGGGGACGCTTACCACG(TGG)) was cloned into the pU6B plasmid as described in (Ren *et al*. 2013). Dany (CG30401) genomic sequences from 1253 bp upstream of the ATG to 307 bp downstream of stop the codon were PCR amplified from Nanos-Cas9 (3^rd^ chromosome) flies (Ren *et al*, 2013) as two fragments and stitched together at the start codon ATG with a SmaI site introduced in the primer. The DNA sequence of V5 (GGTAAGCCTATCCCTAACCCTCTCCTCGGTCTCGATTCTACG) was inserted right after the start ATG in the PCR primer. The resulting 2485 bp *V5-Dany* genomic donor sequence was cloned into the SalI and NotI sites of pBS-SK+ using restriction sites introduced in the PCR primers. Two silent mutations were then introduced to the CRISPR seed sequences in this donor plasmid by site directed mutagenesis to prevent Cas9 cutting of the donor. 200 ng/µl of the pBS_V5-Dany donor plasmid was mixed with 150 ng/µlpU6B plasmid containing the guide sequences and injected into embryos of Nanos-Cas9 flies. Injected flies were allowed to mate to TM3/TM6b; *Sp*/CyO double balancer flies and produce progeny before screening by PCR for genomic insertion of the V5 sequence. 20 CyO virgin female progeny from each PCR positive parent were single-clone mated to double balancer males, and allowed to lay eggs for one week before being PCR screened again for germline clones. Testes from 20 males of each balanced line were dissected and stained with an anti-V5 antibody to confirm expression of V5-Dany specifically in spermatocytes. Positive lines were made homozygous for further characterization.

### Immunofluorescence staining

Testes from 0–2 day old males were dissected in 1× PBS on a cyclops dissecting dish for <30 min at room temperature (RT). Immunostainings were performed following the protocol described by Gallicchio *et al*. 2024. Primary antibodies used were rabbit anti-Kmg (1:10,000 serum AA, used in Kim *et al*. (2017)), chicken anti-GFP (1:10,000; Abcam 13970), and mouse anti-V5 (1:200; Invitrogen 46-0705). Secondary antibodies used were donkey or goat anti-rabbit, anti-mouse, or anti-chicken, all used at 1:500 and conjugated with either Alexa 488, Alexa 568, or Alexa 647 (all from Thermo Fisher). Imaging was performed on a Leica SP8 confocal microscope.

### HCR RNA-FISH

Testes were dissected from 0–2 day old males in 1× PBS on a cyclops dissecting dish for <30 min at room temperature (RT), fixed, and permeabilized as done for immunofluorescence. The protocol for “generic sample in solution” (Rev9) from the Molecular Instruments website (https://www.molecularinstruments.com/hcr-rnafish-protocols) was followed, with alterations to the protocol as described by Gallicchio *et al*. (2024). Probes were designed as in Bedbrook *et al*. 2023 and were obtained from Integrated DNA Technologies as oligo pools (50 pmol/oligo; oPools). Hybridization buffer, wash buffer, amplification buffer, and hairpins (conjugated with either 488 or 546 fluorophores) were obtained from Molecular Instruments (HCR RNA-FISH bundle). Probe sequences are listed in Supplemental Table S1. Imaging was performed on a Leica SP8 confocal microscope.

### Western blots

Testes from 15 0–2 day old males were quickly dissected into 1× PBS in a cyclops dissecting dish at room temperature, then together transferred to a 1.7 ml Eppendorf tube containing 1× PBS. The 1× PBS was then removed, the tube was snap-frozen in liquid nitrogen and stored at −80°C. The western blots were perfomed as described by Gallicchio *et al*. (2024), with the following alterations: lysis buffer used was JUME buffer (10 mM MOPS, 10 mM EDTA, 8 M urea, 1% SDS). Primary antibodies used for Western blots were mouse anti-Taf1 (1:200; 30H9 hybridoma supernatant, a kind gift from Bob Tjian), mouse anti-Chro (1:10,000; 6H11), mouse anti-Hfp (1:200; 6G10-12), rabbit anti-GFP (1:1,000 ; Cell signaling 2956S), rabbit anti-Kmg (1:100,000 serum AA, purified for use in Kim *et al*. (2017)), mouse anti-V5 (1:1,000; Invitrogen 46-0705). Secondary antibodies used for Western blots were peroxidase AffiniPure goat anti-rabbit IgG (H+L; 1:10,000; Jackson ImmunoResearch AB_2313567) and peroxidase AffiniPure goat anti-mouse IgG (H+L; 1:10,000; Jackson ImmunoResearch AB_10015289).

### Immunoprecipitation

Protein A Dynabeads (Thermo Fisher Scientific, #10002D) were used with rabbit anti-Kmg (Kim *et al*. 2017), and pan-mouse IgG Dynabeads (Thermo Fisher Scientific, #11041) were used for anti-GFP (mouse, Millipore Sigma #11814460001). Testes were dissected into 1× PBS in a cyclops dissecting dish in batches of 50 flies for <30 min at room temperature and transferred into a 1.7 ml Eppendorf tube with 1× PBS. The 1× PBS was removed and the tube snap-frozen in liquid nitrogen and stored at −80°C. For immunoprecipitation followed by western blot (Co-IP), 150 pairs of testes per sample were used. Protocol as described by Baker *et al*. (2023) with the following alterations: neither RNAse A, nor SUPERasin were added to the lysis buffer; the supernatant from the IP step was kept to run as a control lane; proteins from Input, IP and supernatant samples were precipitated (protocol below) prior to Western blot (protocol above). For immunoprecipitation followed by mass spectrometry (IP-MS), used protocol as described by Baker *et al*. (2023). 1000 pairs of testes per sample were used. Testes from bamGal4 served as wild type sample (labeled NM_WT_1_171009) and Kmg knockdown testes as a negative control sample (labeled NM_KD_2_171009). Samples were processed and analyzed by Stanford University Mass Spectrometry core. Complete table with IP-MS data in Supplemental Table S2.

### Protein precipitation

To 1 vol of protein sample (in 1.7 ml Eppendorf), 4 vol of methanol (ice-cold) were added. The mixture was vortexed, then 1 vol of chloroform was added. After vortexing again, 3 vol of H2O were added, the mixture was vortexed, then centrifuged at maximum speed for 5 min at 4°C. The upper liquid layer was removed, then 3 vol of ice-cold methanol were added and the mixture was vortexed. The tube was centrifuged at maximum speed for 10 min at 4°C. All liquid was removed, the protein pellet was air-dried and resuspended in 40μl of JUME buffer (10 mM MOPS, 10 mM EDTA, 8 M urea, 1% SDS).

### PacBio library preparation and sequencing

Testes were dissected from 0–2 day old males in 1xPBS, as batches of 50 flies, in a cyclops dissecting dish for <30min at room temperature. Each batch was transferred to a 1.7 ml Eppendorf tube containing 1ml of 1xPBS, the PBS was then immediately removed, and the testes were snap-frozen in liquid nitrogen and stored at −80°C. RNA was extracted from 150 pairs of testes using the RNeasy Plus Mini Kit from QIAGEN (74104). Frozen tissue was dissociated using a 1 ml syringe with a 27-gauge needle aspirating up and down ∼10 times in 300µl of lysis buffer (from RNeasy kit) supplemented with 1:100 β-mercaptoethanol. The quality of extracted RNA was assessed via Bioanalyzer, and the concentration was measured by Qubit RNA HS assay kit (Life Technologies Q32852). Iso-Seq SMRTbell® libraries were made following the “Procedure & Checklist – Iso-Seq™ Express Template Preparation for Sequel® and Sequel II Systems”, protocol from PacBio. Reagents from the following kits were used for library preparation and sequencing: SMRT Bell Express Template express kit 2.0 (100-938-900), Iso-Seq Express Oligo Kit -PacBio (PN 101-737-500), NEBNext® Single Cell/Low Input RNA Library Prep Kit for Illumina (E6420S), Sequel® II Binding Kit 2.1 (101-843-000), Sequel® II Sequencing Kit 2.0 (101-820-200). Each sample was sequenced on 1 SMRT Cell 8M (101-389-001), on a Sequel IIe system (24hr run time) at the Chan Zuckerberg Biohub SF Genomics Platform. Circular consensus sequencing (CCS) and Iso-Seq analysis were performed in SMRT Link v11.0.

### RNA-seq library preparation and sequencing

RNA extraction was done as for PacBio samples, from 150 pairs of testes. Library preparation was carried out using the NEBNext® Ultra™ II Directional RNA Library Prep Kit for Illumina (#E7760S), with the NEBNext® Poly(A) mRNA Magnetic Isolation Module (#E7490S). Two biological replicates were performed per condition. Sequencing was performed by Novogene on a NOVAseq, PE 150 Illumina platform.

### 3′ end library preparation and sequencing

RNA extraction was done as for PacBio samples, from 150 pairs of testes. Library preparation was carried out using the Quant Seq (FWD) kit from Lexogen. Two biological replicates were performed per condition. Sequencing was performed by the Stanford University Genomics Facility on an Illumina NextSeq 500 platform, as in (Berry *et al*. 2022).

### CAGE library preparation and sequencing

RNA extraction was done as for PacBio samples. 10ug of total RNA were sent to DNAform (https://www.dnaform.jp/en/products/library/cage/) for CAGE library preparation using published protocol nAnTi-CAGE (non-amplifying-non-tagging illumina CAGE) (Murata *et al*. 2014). Control samples (bamGal4) were sequenced using Illumina Next-seq, with 150bp PE sequencing (only read1 was used for analysis). Kmg knockdown (KD) samples were sequenced using Illumina HiSeq with 75-bp SE sequencing. Two biological replicates were performed per condition.

### ATAC-seq library preparation and sequencing

ATAC-seq was carried out with a modified version of the following published protocol (Buenrostro *et al*. 2015), as described by Lu *et al*. (2020). Two biological replicates were performed per condition. Sequencing was done with HiSeq 4000 at the Stanford Functional Genomics Facility, with 75bp PE.

### ChIP-seq library preparation and sequencing

ChIP-seq was performed using the Low Cell ChIP Kit from Active Motif (#53086). Testes from 0–2 day old males were dissected in 1xPBS, in batches of 50-100 flies, in a cyclops dissecting dish for <30min at room temperature. Each batch was transferred to a 1.7ml Eppendorf tube containing 1ml of 1xPBS, then immediately fixed by adding 28µl of 37% Formaldehyde (Sigma-Aldrich #252549), and the sample incubated 10min at RT with rotation. The reaction was quenched by adding 52.6µl of glycine (2.5M) and incubated 5min at RT. The liquid was removed and 1ml of 1xPBS was added. After repeating this wash the PBS was removed, the tube was snap-frozen in liquid nitrogen and stored at −80°C. Protein G Agarose Beads were prepared following “Session D” of the Low Cell ChIP Kit protocol. While incubating the “pre-clearing” beads, the chromatin sample was prepared as follow: pool 350 pairs of frozen testes in 130µl of ChIP buffer mix (from Low Cell ChIP Kit); the tissue was dissociated using a 1 ml syringe with a 27-gauge needle aspirating up and down ∼10 times; the sample was transferred to microTUBE-130 (microTUBE AFA Fiber Pre-Slit Snap-Cap, #520045), kept on ice, and sonicated using Covaris (E220 Evolution) (treatment: 600sec; Peak Power: 75; duty factor: 10; cycles/burst: 200; avg power: 17.5, used intensifier); the sample was then transferred to new a 1.7 ml Eppendorf tube, centrifuged 5min at 4°C, and the supernatant transferred to new tube; the volume of the transferred sample was brought to 200µl by adding ∼70µlof ChIP buffer mix; 5% of input (10ul) was removed and saved at 4°C until needed. Pre-clearing of the Chromatin, Immunoprecipitation, and Washing & Elution of IP reactions were done as described in “Session E, F and G” of the Low Cell ChIP Kit protocol. Antibodies used for the Antibody Mixture: rabbit anti-Kmg (Kim *et al*. 2017), mouse anti-V5 (Invitrogen 46-0705), or rabbit anti-HA (Cell Signaling C29F4). The eluted ChIP DNA (end of section G) was transferred to a 250µl PCR tube, and 2µl of Proteinase K and 5µl of 5 M NaCl were added. Tubes were vortexed to mix and kept at a 65°C overnight, in a thermocycler, to reverse cross-links. For the input samples: 90µl of TE and 2µl of RNase A were added and the mixture was incubated for 30min and 37°C. Then the input samples were transferred to PCR tubes, 2µl of Proteinase K and 5µl of 5 M NaCl were added and the mixtures incubate over-night at 65°C. An alternative DNA purification protocol, to that proposed by the kit manufactures, was used: samples and input samples were moved to new 1.7 ml Eppendorf tubes, 1.8x volume of SPRIselect beads (Beckman Coulter #B23318) were added, mixed by pipetting and incubated 15 min at RT. The beads were magnetically separated, washed 2x with 80% Ethanol, and DNA was eluted with 50µl of TE. Libraries were made using NEBNext® Ultra™ II DNA Library Prep Kit for Illumina® NEB (#E7645L). Notes on protocol: Due to low concentration, input DNA was not quantified, instead all 50ul of eluted DNA was used. Adaptor was diluted 1:25. We skipped “3A Size Selection of Adaptor-ligated DNA”. Two biological replicates were performed per condition. Sequencing was performed by Novogene on a NOVAseq, PE 150, Illumina platform.

### PacBio data analysis

Processing of each library of PacBio HiFi reads was done following the IsoSeq pipeline (ccs, lima, IsoSeq (3.4.0)). Data were mapped to *Drosophila* genome assembly: BDGP6.32 (GCA_000001215.4) using pbmm2 (1.12.0). Tama (b0.0.0) collapse was used to collapse redundant transcript models (Kuo *et al*. 2020); -x capped (although our libraries are not 5’ capped, this avoids over collapsing), -a 78 (78bp interquantile width that includes 90% of CAGE clusters, from parallel analysis), -z 80 (80bp interquantile width that includes 90% of 3’end clusters, from parallel analysis). See analysis pipeline in Supplemental figure 1.

### RNA-seq data analysis

RNA-seq data were used independently in 2 pipelines. 1) Used as input files to run SQANTI3, which used Kallisto to map reads to transcriptome and quantify transcript isoforms (details below). 2) Adapters and low-quality bases were trimmed with trimGalore (0.6.6) (Martin 2011); reads were mapped to *Drosophila* genome assembly: BDGP6.32 (GCA_000001215.4) using STAR (2.7.9a) (Dobin *et al*. 2013); DeepTools (3.3.0) was used to generate normalized coverage tracks for visualization and plot heatmaps (Ramírez *et al*. 2016).

### CAGE data analysis

Low-quality bases were trimmed using TrimGalore (0.6.6) (Martin 2011) and mapped to *Drosophila* genome assembly: BDGP6.32 (GCA_000001215.4) using STAR (2.7.11b) (Dobin *et al*. 2013). The most 5′ nucleotide of each mapped read was counted using bedtools coverage (2.27.1) (Quinlan and Hall 2010) and used as input into CAGEr to build CAGE clusters (1.20.0) (Haberle *et al*. 2015).

### 3’ end seq data analysis

3’ end raw data for control samples (bamGal4) used here were published in Gallicchio *et al*. (2024), genomic data available under NCBI GEO (GSE272579). Kmg knockdown samples were generated for this publication. Low-quality bases, polyA tails and adapters were trimmed using bbmap (39.01) and reads were mapped to *Drosophila* genome assembly: BDGP6.32 (GCA_000001215.4) using STAR (2.7.11b). The most 3′ nucleotide of each mapped read was counted using bedtools coverage (2.27.1) and used as input into CAGEr to build 3’ end clusters (1.20.0).

### ATAC-seq data analysis

ATAC-seq raw data for control (ctrl^72hrs^), and *aly* mutant samples used here were published in Lu *et al*. (2020), genomic data available under NCBI GEO (GSE145975). Kmg knockdown samples were generated for this publication. Analysis done exactly as described by Lu *et al*. (2020).

### ChIP-seq data analysis

ChIP-seq data for HA-Aly and Mi-2 samples used here were published in Kim *et al*. (2017), raw data available under NCBI GEO (GSE89506). All other samples were generated for this publication. Adapters and low-quality bases were trimmed using trimGalore (0.6.6), reads were mapped to *Drosophila* genome assembly: BDGP6.32 (GCA_000001215.4) using bwa aln (0.7.18) (Li and Durbin 2009). DeepTools (3.1.0) was used to generate normalized coverage tracks for visualization and plot heatmaps. MACS2 (2.1.1) was used for peak calling (Zhang *et al*. 2008).

### Build and quantify transcriptome

Transcriptome models from each PacBio library were merged into one single transcriptome with Tama (b0.0.0) merge (Kuo *et al*. 2020); -a 78 (78bp interquantile width that includes 90% of CAGE clusters, from parallel analysis), -z 80 (80bp interquantile width that includes 90% of 3’end clusters, from parallel analysis). SQANTI3 was used to integrate our transcriptome model with the *Drosophila* annotation BDGP6.32.109, predict ORF for each transcript isoform, map short-reads to the transcriptome model and predict RT-switching artifacts (Pardo-Palacios *et al*. 2024). To focus on full length transcripts, the transcriptome model was then filtered to keep only transcripts that met the following criteria: their TSS overlapped with CAGE consensus clusters, their TTS overlapped with 3’ end consensus clusters, all splice-junctions had RNA-seq coverage and the transcript had no RT switching artifacts. To quantify transcript isoform expression we used RNA-seq data to run SQANTI3 (with Kallisto). tappAS was used for transcript expression analysis, and expression of predicted protein isoforms (Fuente *et al*. 2020). Check analysis pipeline in Supplemental Figure 1.

### Quantification of promoter output

To define promoters, coordinates of the transcription start site (TSS) for each transcript isoform were intersected with coordinates of CAGE consensus clusters (from CAGE analysis), using bedtools (2.27.1). The ID of the CAGE consensus cluster was considered the promoter ID. To quantify expression from a given promoter, the expression from all transcripts starting at that promoter was summed. Used DEseq2 for differential promoter expression analysis (Love, Huber, and Anders 2014).

### Definition of cryptic promoter and cryptic protein

Cryptic promoters were defined as having an expression level in Kmg knockdown sample (from promoter output analysis) higher than quantile 0.25, being upregulated 16 fold or more in Kmg KD compared to control; having a padj value lower than 0.05; and the promoter (defined by CAGE cluster coordinates) did not contain any annotated TSS (from BDGP6.32). Cryptic proteins were defined by deriving from cryptic promoters, having an expression level in Kmg knockdown sample (from tappAS analysis) higher than quantile 0.25, upregulated 16 fold or more in Kmg knockdown compared to control, having a padj value lower than 0.05.

### Analysis of Aly ChIP-seq peaks

To define one TSS per Aly peak, we intersected Aly ChIP-seq peak coordinates with promoter coordinates using bedtools. We selected the highest expressed promoter that overlaps with each Aly peak. From the selected promoter we chose the highest expressed transcript and got its TSS coordinate, which was used for downstream analysis (below). Classification of Aly peaks, relative to which type of promoter Aly binds to (Aly-dependent, Aly-independent, or cryptic promoter), was done based on the highest expressed promoter it overlapped with.

### Downstream analysis

DeepTools was used to generate matrixes used for graphs of enrichment profiles, and to plot heatmaps (3.3.0) (Ramírez *et al*. 2016). Graphs were generated using ggplot2 (Wickham 2016). Integrative Genomics Viewer (IGV) was used for genomic data visualization (Robinson *et al*. 2011). SRplot was used to plot the chromosome distribution of cryptic promoters (Tang *et al*. 2023).

## Data availability

All sequencing data were submitted to GEO (GSE…). Analysis scripts are available at https://github.com/NeuzaRM/Kmg_FullerPaper2025. Analysis script for cryptic protein classification is available at https://github.com/julianignacioperez/Kmg_FullerPaper2025.

## Acknowledgments

We thank all members of the Fuller laboratory for feedback, and Dr. Tomek Swigut for constructive discussions. We thank Dr. Lauren Goins for feedback and critical reading of the manuscript. Special thanks to Norma Neff and her team at CZ Biohub SF for supporting all the PacBio long read sequencing. We thank Dr. Bob Tjian for Taf1 antibody. We thank the Stanford Fly Media Center and all the staff working in the Department of Developmental Biology. We also thank the Stanford Cell Science Imaging Facility (CSIF), Protein and Nucleic Acid Facility (PAN) and Vincent Coates Foundation Mass Spectrometry Laboratory, and the Stanford University Mass Spectrometry core (SUMS). We extend a big thank you to the Bloomington Drosophila Stock Center, Vienna Drosophila Resource Center and the Drosophila Genomics Resource Center for making fly strains readily available. Finally, we would like to acknowledge FlyBase as an indispensable resource. N.R.M was supported by an EMBO Long-Term Postdoctoral Fellowship (ALTF 1551-2015). This research was supported by NIH grant R35 GM136433 and funds from the Katherine D. McCormick and Stanley McCormick Memorial Professorship, and the Reed-Hodgson Professor ship in Human Biology to M.T.F.

## Author Contributions

N.R.M. and M.T.F. conceived the project and wrote the manuscript. N.R.M. designed the experimental work and performed or supervised all experimental and computational procedures (except ATAC-seq). L.G. carried out immunofluorescence, RNA FISH and Western Blot experiments. D.L. carried out ATAC-seq experiments and ATAC-seq data analysis. J.J.K. generated preliminary microarray analysis on Dany knockdown (not published). J.P. carried out analysis of cryptic protein classification. A.M.D. carried out PacBio library sequencing. C.L. generated the V5-*Dany* CRISPR line. B.B helped with *Drosophila* strain construction. Funding was acquired by M.T.F.

